# The “don’t eat me” signal CD47 contributes to microglial phagocytosis defects and autism-like behaviors in 16p11.2 deletion mice

**DOI:** 10.1101/2024.06.07.597763

**Authors:** Jun Ju, Yifan Pan, Xinyi Yang, Xuanyi Li, Jinghong Chen, Shiyu Wu, Sheng-Tao Hou

## Abstract

Various pathological characteristics of autism spectrum disorder (ASD) stem from abnormalities in brain resident immune cells, specifically microglia, to prune unnecessary synapses or neural connections during early development. Animal models of ASD exhibit an abundance of synapses in different brain regions, which is strongly linked to the appearance of ASD behaviors. Overexpression of CD47 on neurons acts as a “don’t eat me” signal, safeguarding synapses from inappropriate pruning by microglia. Indeed, CD47 overexpression occurs in 16p11.2 deletion carriers, causing decreased synaptic phagocytosis and the manifestation of ASD characteristics. However, the role of CD47 in synaptic pruning impairment leading to ASD phenotypes in the 16p11.2 deletion mouse model is unclear. Moreover, whether blocking CD47 can alleviate ASD mice’s behavioral deficits remains unknown. Here, we demonstrate a strong link between increased CD47 expression, decreased microglia phagocytosis capacity, and increased impairment in social novelty preference in the 16p11.2 deletion mice. The reduction in microglia phagocytosis caused a rise in excitatory synapses and transmission in the prefrontal cortex of 16p11.2 deletion mice. Importantly, blocking CD47 using a specific CD47 antibody or reducing CD47 expression using a specific shRNA enhanced the microglia phagocytosis and reduced excitatory transmission. Reduction in CD47 expression improved social novelty preference deficits in 16p11.2 mice. These findings demonstrate that CD47 contributes to the ASD phenotypes in the 16p11.2 deletion mice and could be a promising target for the development of treatment for ASD.

**Significance Statement:** Autism spectrum disorder (ASD) is a neurological developmental condition characterized by stereotyped behaviors and cognitive deficits. However, therapeutic options for ASD remain limited. Activation of the classical complement system, an innate immune signaling pathway component, supports microglia-mediated synaptic pruning during development and disease. In particular, CD47, a “don’t eat me” signal, protects synapses from inappropriate clearance. Here, we investigated the role of CD47 in microglial phagocytosis using the 16p11.2 deletion mouse model, demonstrating that reducing CD47 signaling enhances microglial phagocytose synapses in the prefrontal cortex. This enhancement leads to improved synaptic function and reduced social behavioral deficits. These findings provide mechanistic insights into the role of CD47, laying the groundwork for developing more effective treatments for ASD.

## Introduction

Copy number variation at the chromosomal 16p11.2 locus is strongly linked to autism spectrum disorder (ASD) (1–3), a developmental neurological condition characterized by stereotyped behaviors and cognitive deficits (4–6). The 16p11.2 deletion mouse carries a deletion of the human equivalent of the 16p11.2 region and displays behavioral abnormalities similar to those found in individuals with the 16p11.2 chromosomal deletion syndrome (2, 3). Thus, the 16p11.2 deletion mouse model is a valuable tool for studying the genetic and behavioral mechanisms of ASD and serves as a platform for developing potential therapeutic interventions.

Various pathological characteristics of ASD stem from functional abnormalities of brain resident immune cells, particularly microglia (7–9). Microglia regulate synaptic circuit remodeling and phagocytose synaptic material in healthy and diseased human and mouse brains (8, 10–13). During development, microglia engulf neuronal precursors and contribute to synaptic pruning mechanisms (11). In the adult brain, microglia play a role in influencing synaptic signaling and shaping synaptic plasticity. In fact, evidence suggests that the maturation and function of distinct neural circuits may potentially be linked to the molecular identity that microglia adopt across the brain (10, 14–16). Our recent studies using the 16p11.2 deletion mouse model demonstrated a significant reduction in microglia phagocytosis, enhanced excessive excitatory neurotransmission of the pyramidal neurons in the prefrontal cortex (PFC), and their linkage with social novelty deficit (17). However, the molecular mediators for microglial regulation of synaptic steady-state in the 16p11.2 deletion mouse brain remain unclear.

Activation of the classical complement system, part of the innate immune signaling pathway, supports microglia-mediated synaptic pruning during development and disease (14–16, 18, 19). In particular, CD47, known as a “don’t eat me” signal, protects synapses from inappropriate clearance (14, 18). Overexpression of CD47 in neurons and loss of its microglial receptor, SIRPα, inhibit microglia engulfment or phagocytosis, thus being implicated in the development of various neurological diseases such as Alzheimer’s disease (AD) (15) and ASD (18). CD47 is localized on active synapses, suggesting that synaptic pruning is activity-dependent and that CD47 may protect the highly active synapses from microglial pruning (14). Knockdown of CD47 or SIRPα leads to failure to display preferential engulfment of less active inputs and over-pruning synapses during postnatal development (14), in AD brain (15), and axonal degenerative conditions (20).

The PFC is a crucial brain region regulating social behaviors and is closely associated with ASD. Individuals with 16p11.2 deletion exhibit impaired PFC connectivity with other brain regions involved in sociability functions (21). Immunohistochemical studies demonstrated elevated levels of dendritic spines in the 16p11.2 deletion mouse PFC, and electrophysiological data showed that 16p11.2 deletion mice have decreased PFC neuronal activity and abnormal NMDA receptor function (1, 17, 22). Chemogenetic activation of PFC could rescue synaptic and behavioral deficits in the mouse model of 16p11.2 deletion syndrome (22).

However, the specific role of CD47 in the 16p11.2 deletion mouse brain, particularly its impact on synaptic pruning impairment that contributes to ASD phenotypes, remains uninvestigated. In addition, it is also unclear whether blocking CD47 can effectively alleviate ASD behavioral deficits. In this study, we examined the influence of CD47 on microglia phagocytosis in the 16p11.2 deletion mice PFC and demonstrated that inhibiting CD47 signaling enhances microglia’s capacity to phagocytose synapses. This enhancement leads to improved synaptic function and a reduction in social novelty deficits.

## Results

### Impairment in recognition memory and social novelty ability in 16p11.2 deletion mice

Several behavioral tests were performed on the 16p11.2 deletion mice (labeled 16p in all figure panels). In the open field test, behavioral parameters, such as distance traveled, movement velocity, and time spent in the open field center, were unaffected, indicating similar locomotion and anxiety levels in both 16p11.2 deletion mice and the wildtype littermates (WT) (Fig. 1 A–D). The alternation index also remained unchanged in the Y maze test (Fig. 1 E and F). However, in the novel object recognition (NOR) test, the recognition index was significantly reduced, suggesting impaired recognition memory in 16p11.2 deletion mice (Fig. 1 G and H). Furthermore, the 16p11.2 deletion mice exhibited significantly impaired social novelty ability, albeit a normal social ability index during the three-chamber test (Fig. 1 I–K). Collectively, 16p11.2 deletion mice displayed normal locomotion, anxiety levels, and working memory but exhibited deficits in recognition memory and social novelty ability.

**Figure 1.**
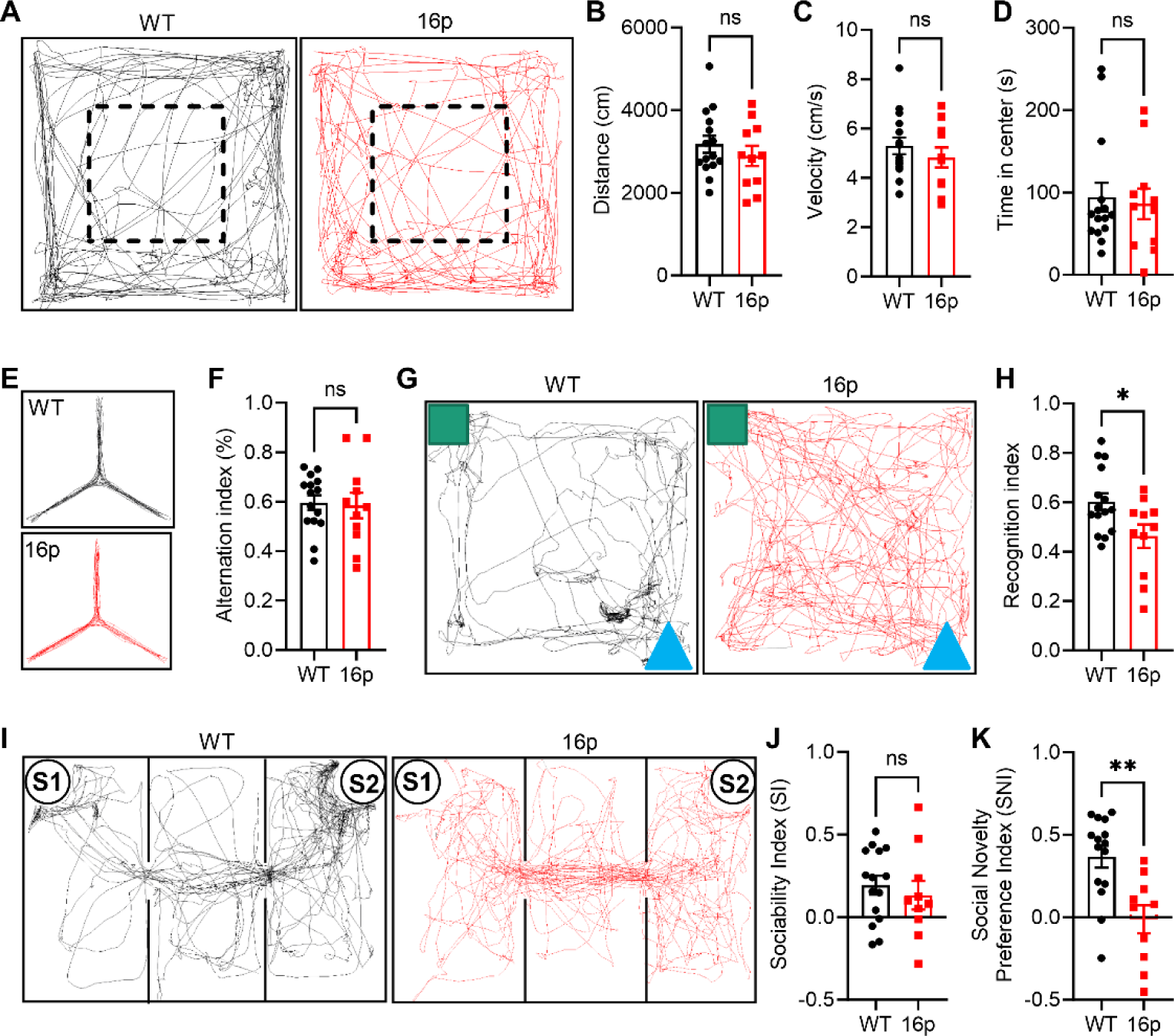
Determination of recognition memory and social novelty deficits in 16p11.2 deletion mice. A The trajectory chart of mice in the open field. The dash-lined box: center area. B The distance traveled in the open field. C The velocity traveled in the open field. D Time in the center in the open field. E The trajectory chart of mice in Y maze. F Alternation index in the Y maze. G The trajectory chart of mice in novel object recognition test (NOR). Green square: familiar object; Blue triangle: novel object. H Recognition index in the NOR test. I The trajectory chart of mice in the three-chamber test. S1: familiar mouse, S2: novel mouse. J Sociability index in the three-chamber test. K Social novelty preference index in the three-chamber test. ns, not significant, *P < 0.05, **P < 0.01, unpaired *t*-test for panels B, C, F, H, J and Mann-Whitney U test for panels D, K for analysis. The numbers of mice used for experiments are shown in A – I: n = 15 (WT) and n = 11 (16p). For panel K, WT: n = 15 mice, 16p: n = 10 mice.

### Reduced microglial phagocytosis in 16p11.2 deletion mice

To examine microglia alterations in 16p11.2 deletion mice, we employed western blotting to assess expressions of crucial marker proteins (Fig. 2 A). The expression level of CORO1A, a gene located within the 16p11.2 deletion region, was significantly reduced, confirming the fidelity of the 16p11.2 deletion mouse model (Fig. 2 A and B). Furthermore, the expression level of IBA1, a microglial biomarker, was significantly reduced in the 16p11.2 deletion mice (Fig. 2 A and C). To further substantiate IBA1 change in microglia, we quantified IBA1 immunostaining density across various brain regions, including the PFC, primary motor cortex, sensory cortex, and striatum. The result revealed no significant difference in microglia density in these brain regions between the 16p11.2 deletion mice and the WT littermates (Fig. 2 D and E, SI Appendix, Fig. S1).

**Figure 2.**
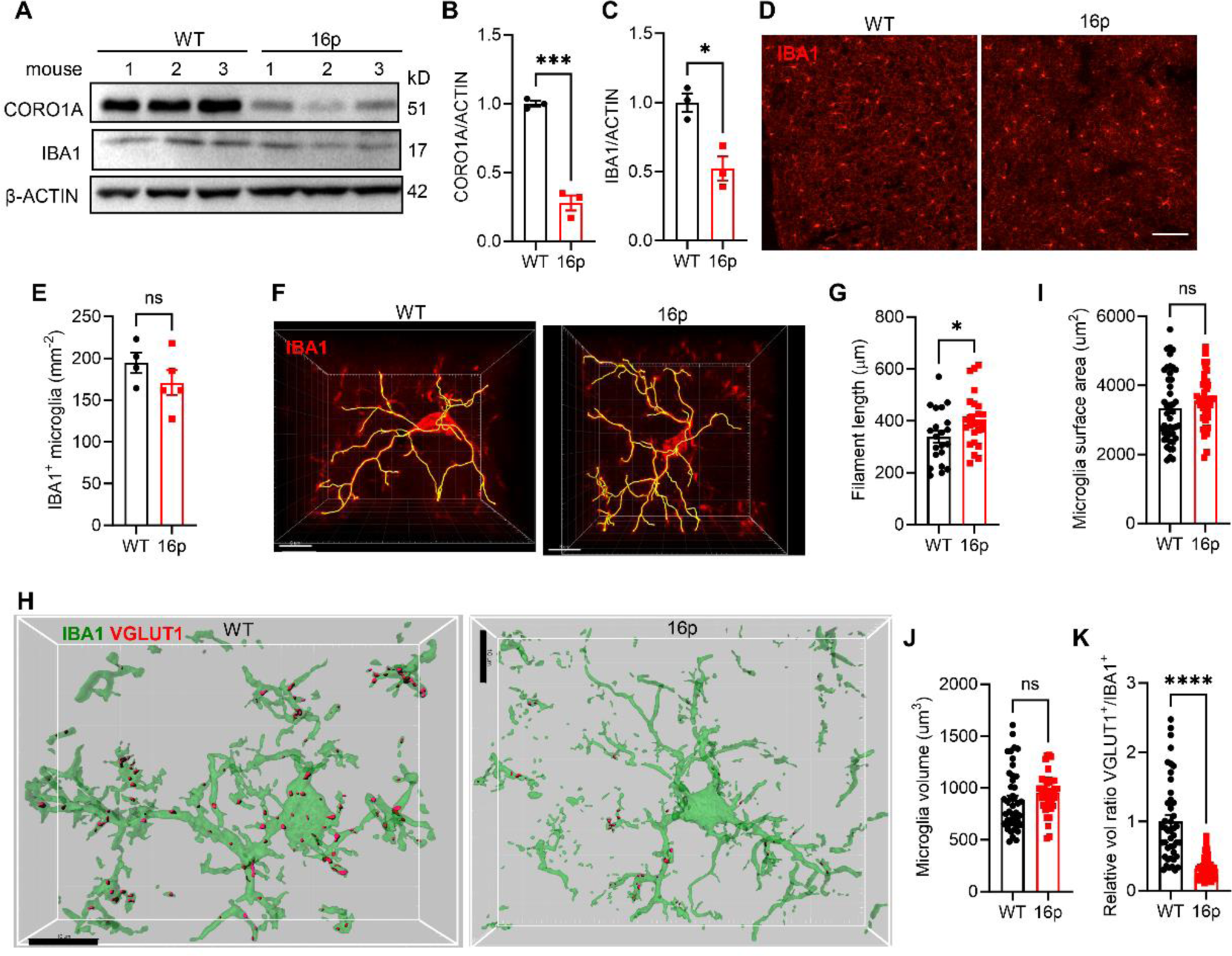
Reduction of microglia-dependent synapse pruning in 16p11.2 deletion mice. A Western blotting images of CORO1A and IBA1 in WT and 16p group of mice. The expression of β-ACTIN was used as an internal loading control. B Quantification of CORO1A expression in PFC. C Quantification of IBA1 expression in PFC (WT: n = 3 mice, 16p: n = 3 mice). D Representative images of IBA1 immunostaining in WT and 16p mice PFC. The scale bar: 100 μm. E Quantification of IBA1^+^ microglia density (WT: n = 4 mice, 16p: n = 5 mice). F Representative images of a single microglia. The scale bar: 10 μm. G Quantification of filament length of individual microglia cells (WT: n = 21 cells from 4 mice, 16p: n = 25 cells from 5 mice). H Representative images of IBA1 and VGLUT1 co-immunostaining in WT and 16p mice groups. The scale bar: 10 μm. I Quantification of microglia surface area. J Quantification of microglia volume. K Quantification of VGLUT1 volume ratio in microglia (WT: n = 44 cells from 4 mice, 16p: n = 41 cells from 4 mice). ns, not significant, *P < 0.05, ***P < 0.001, ****P < 0.0001, unpaired *t* test for panels B, C, E, G and Mann-Whitney U test for panels I, J, K.

However, when the morphology of microglia from the PFC region was examined using confocal microscopy, we found a significant increase in the ramification of microglial processes in 16p11.2 deletion mice (Fig. 2 F and G), indicating an increase in the resting morphology of microglia. Indeed, the microglia activation marker CD68 expression level was decreased in the 16p11.2 mouse PFC, supporting reduced microglial activation (SI Appendix, Fig. S2). Importantly, the microglial phagocytosis was significantly reduced as determined using IBA1 and VGLUT1 co-immunostaining. While the surface area and volume of microglia remained unchanged, the engulfed VGLUT1 in microglia was significantly reduced in 16p11.2 deletion mice compared with the WT littermates (Fig. 2 H–K). These data demonstrated reduced microglia activation and phagocytosis capacity in 16p11.2 deletion mice.

To further demonstrate microglia alternation in 16p11.2 deletion mice, we sought to activate microglia by administering lipopolysaccharide (LPS). Interestingly, a significant enhancement in the engulfment of synapses occurred in 16p11.2 deletion mice after 7 days of LPS treatment (SI Appendix, Fig. S3). Together, these findings demonstrated that microglia in 16p11.2 deletion mice were at a reduced activation and phagocytose state, which might contribute to the lack of adequate clearance of excessive synapses in ASD brain.

### Increased excitatory synapse number and synaptic transmission in 16p11.2 deletion mice

Experiments were designed to determine whether altered microglial phagocytosis affects the quantity and function of excitatory synapses. Using Golgi staining, we observed a significant increase in dendritic spine numbers in 16p11.2 deletion mice, indicating a defect in microglia clearance (Fig. 3 A and B). Electrophysiological measurement of spontaneous excitatory postsynaptic currents (sEPSCs) in the PFC of 16p11.2 deletion mouse showed an increased sEPSC frequency, but not amplitude, compared with the WT littermates (Fig. 3 C-E). The paired-pulse ratio (PPR) in 16p11.2 deletion mice was also reduced, indicating an elevated release probability of excitatory synapses in 16p11.2 deletion mice (Fig. 3 F and G).

**Figure 3.**
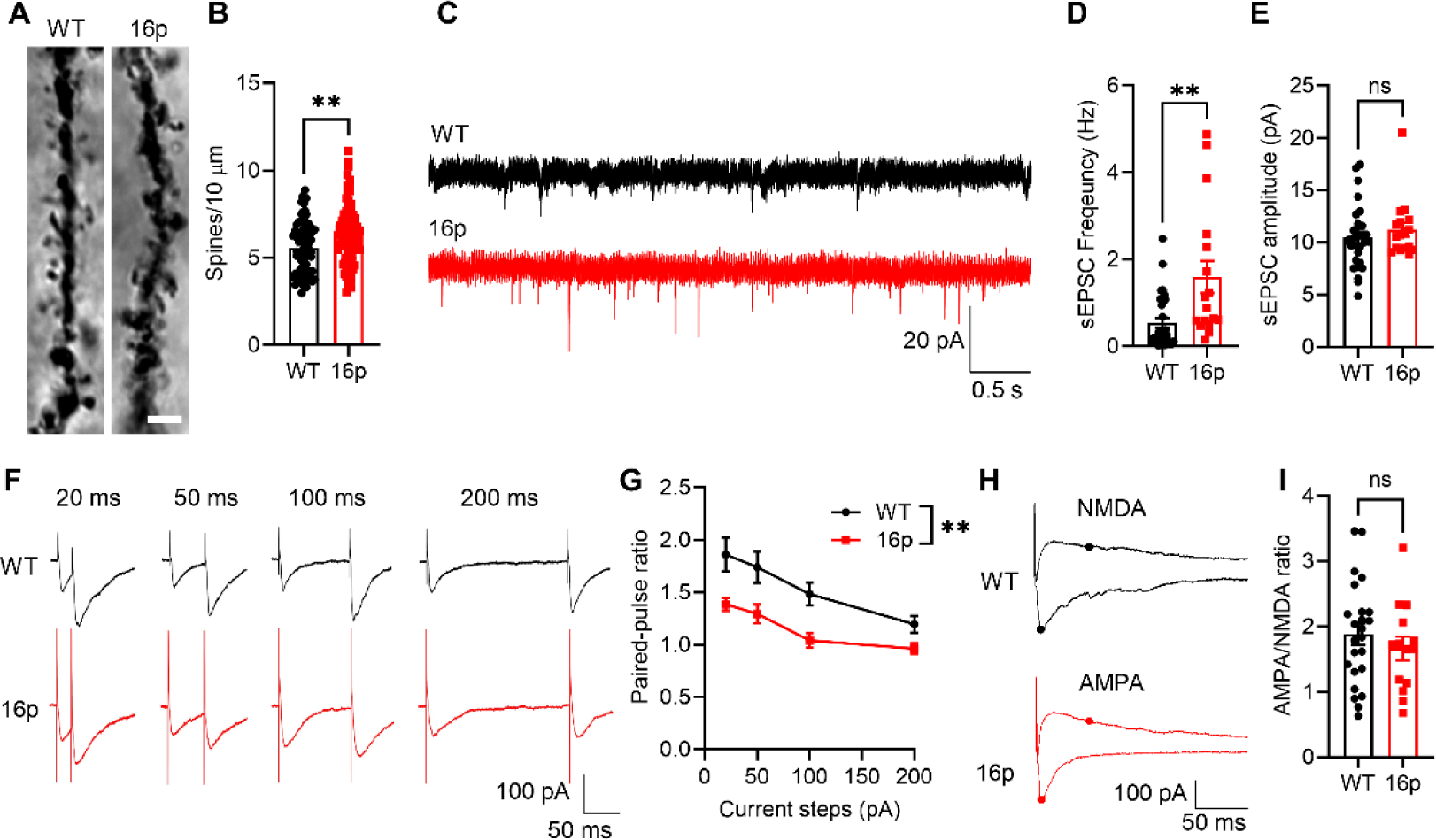
Increased excitatory synapses and excitatory transmission in 16p11.2 deletion mice. A Representative images of Golgi staining of WT and 16p mice, scale bar = 3 μm. B Quantification of numbers of dendritic spines (WT: n = 72 dendrites from 3 mice, 16p: n = 83 dendrites from 3 mice). C Representative traces of sEPSC recorded in the PFC of WT and 16p mice. The scale bar: 20 pA/0.5 s. D Quantification of sEPSC frequency. E Quantification of sEPSC amplitude (WT: n = 30 cells from 6 mice, 16p: n = 17 cells from 6 mice). F Representative traces of paired-pulse stimulation at the interval of 20 ms, 50 ms, 100 ms, and 200 ms in WT and 16p mice. The scale bar: 100 pA/50 ms. G Quantification of evoked EPSCs PPR (WT: n = 14 cells from 4 mice, 16p: n = 15 cells from 6 mice). H Representative traces of AMPA/NMDA ratio in WT and 16p mice. The scale bar: 100 pA/50 ms. G Quantification of AMPA/NMDA ratio (WT: n = 24 cells from 6 mice, 16p: n = 14 cells from 6 mice). ns, not significant, **P < 0.01, unpaired *t* test for panels B, I, Mann-Whitney U test for panels D, E and two-way RM ANOVA with Sidak’s multiple comparisons *post hoc* test for panel G.

However, the 16p11.2 deletion mouse PFC neurons exhibit neither an altered ratio of AMPA/NMDA receptors (Fig. 3 H and I), nor changes in the numbers of NeuN-positive mature neurons and parvalbumin-positive (PV) interneurons in the PFC (SI Appendix, Fig. S4). Therefore, these data support the hypothesis that the 16p11.2 deletion mouse PFC has microglial phagocytosis defects, causing increased excitatory synapses and synaptic transmission.

### Anti-CD47 antibody treatment enhanced microglia synaptic pruning and reduced synaptic transmission in PFC of 16p11.2 deletion mice

The neuronal expression of CD47, a negative regulator of phagocytosis, is elevated in individuals with 16p11.2 deletions (18). To confirm the upregulation of CD47 expression in the brain of 16p11.2 deletion mice, we quantified CD47 protein levels using western blotting and observed a significant increase in CD47 in the PFC (Fig. 4 A and B). Immunostaining also confirmed the significantly increased CD47 expression in the PFC of 16p11.2 deletion mice (Fig. 4 C and D). These findings led us to hypothesize that the increased CD47 expression in neurons might impede the phagocytic capacity of microglia.

**Figure 4.**
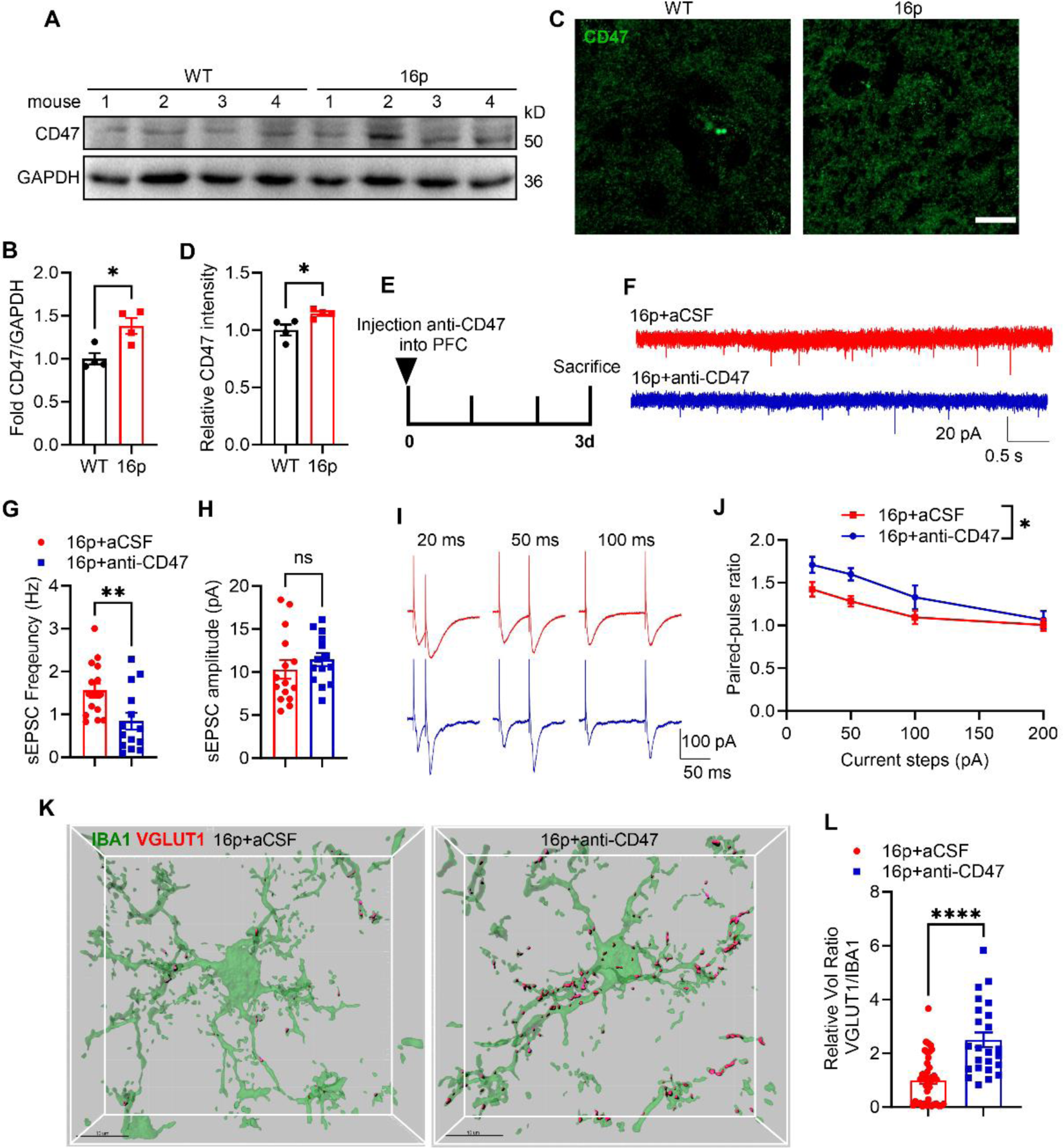
Blocking CD47 using a specific antibody *in vivo* reduced excitatory transmission in 16p11.2 deletion mice. A Western blotting images of PFC CD47 expression levels in the WT and 16p mice. The expression of GAPDH was used as an internal control. B Quantification of CD47 expression (WT: n = 4 mice, 16p: n = 4 mice). C Representative images of PFC CD47 staining in WT and 16p mice. D Quantification of CD47 intensity in PFC (WT: n = 4 mice, 16p: n = 4 mice). E The experimental scheme of anti-CD47 antibody treatment. F Representative traces of sEPSC recorded in PFC. The scale bar: 20 pA/0.5 s. G Quantification of sEPSC frequency. H Quantification of sEPSC amplitude (16p+aCSF: n = 15 cells from 3 mice, 16p+anti-CD47: n = 14 cells from 3 mice). I Representative traces of paired-pulse stimulation at the intervals of 20 ms, 50 ms, and 100 ms in two groups. The scale bar: 100 pA/50 ms. J Quantification of evoked EPSCs PPR (16p+aCSF: n = 14 cells from 3 mice, 16p+anti-CD47: n = 8 cells from 3 mice). K Representative images of IBA1 and VGLUT1 co-immunostaining in two groups. The scale bar: 10 μm. L Quantification of VGLUT1/IBA1 relative volume ratios in microglia (16p+aCSF: n = 41 cells from 3 mice, 16p+anti-CD47: n = 24 cells from 3 mice). ns, not significant, *P < 0.05, **P < 0.01, ****P < 0.0001, unpaired *t* test for panels B, C, H, Mann-Whitney U test for panels G, L and two-way RM ANOVA with Sidak’s multiple comparisons *post hoc* test for panel J.

To test this hypothesis, we investigated the *in vitro* and *in vivo* effects of blocking CD47 using a specific anti-CD47 antibody. Firstly, brain slice preparations from 16p11.2 deletion mice were incubated with the anti-CD47 antibody in artificial cerebrospinal fluid (aCSF). Subsequent measurements of sEPSCs did not reveal any changes in sEPSC frequency and amplitude (SI Appendix, Fig. S5). Secondly, we stereotactically injected the anti-CD47 antibody into the PFC and observed a significant reduction in sEPSC frequency on brain slice preparations after 3 days of anti-CD47 injection. In contrast, the sEPSC amplitude remained unaltered (Fig. 4 E-H). The PPR of evoked EPSCs also significantly increased (Fig. 4 I and J), suggesting a reduced presynaptic release capacity in 16p11.2 deletion mice following anti-CD47 treatment. Finally, co-immunostaining of IBA1 and VGLUT1 revealed enhanced engulfment of VGLUT1 by microglia 3 days after anti-CD47 treatment (Fig. 4 K and L). These findings demonstrated that blocking CD47 enhances synaptic pruning and synaptic transmission in the 16p11.2 deletion mice.

### Reducing CD47 expression via shRNA enhanced microglia synaptic pruning and diminished synaptic transmission in the PFC of 16p11.2 deletion mice

To further confirm the influence of CD47 on PFC neuronal excitability, microglia synaptic pruning, and synaptic transmission, a recombinant adeno-associated virus (rAAV) carrying CD47 shRNA (labeled CD47-shRNA in figure panels) was stereotaxically injected into the PFC (Fig. 5 A). After 3 weeks, we quantified CD47 protein expression levels using western blotting to confirm a significant knockdown of CD47 expression in the PFC by the rAAV CD47-shRNA treatment (Fig. 5 B and C). Immunostaining showed a reduced CD47 signal in the PFC of 16p11.2 deletion mice (Fig. 5 D and E). Electrophysiological recordings showed reduced sEPSC frequency, while the sEPSC amplitude remained unchanged with rAAV CD47-shRNA treatment (Fig. 5 F-H). The PPR of evoked EPSCs was increased, indicating diminished presynaptic release capacity in 16p11.2 deletion mice following rAAV CD47-shRNA treatment (Fig. 5 I and J). Importantly, co-immunostaining of IBA1 and VGLUT1 revealed enhanced engulfment of VGLUT1 by microglia after rAAV CD47-shRNA treatment (Fig. 5 K and L).

**Figure 5.**
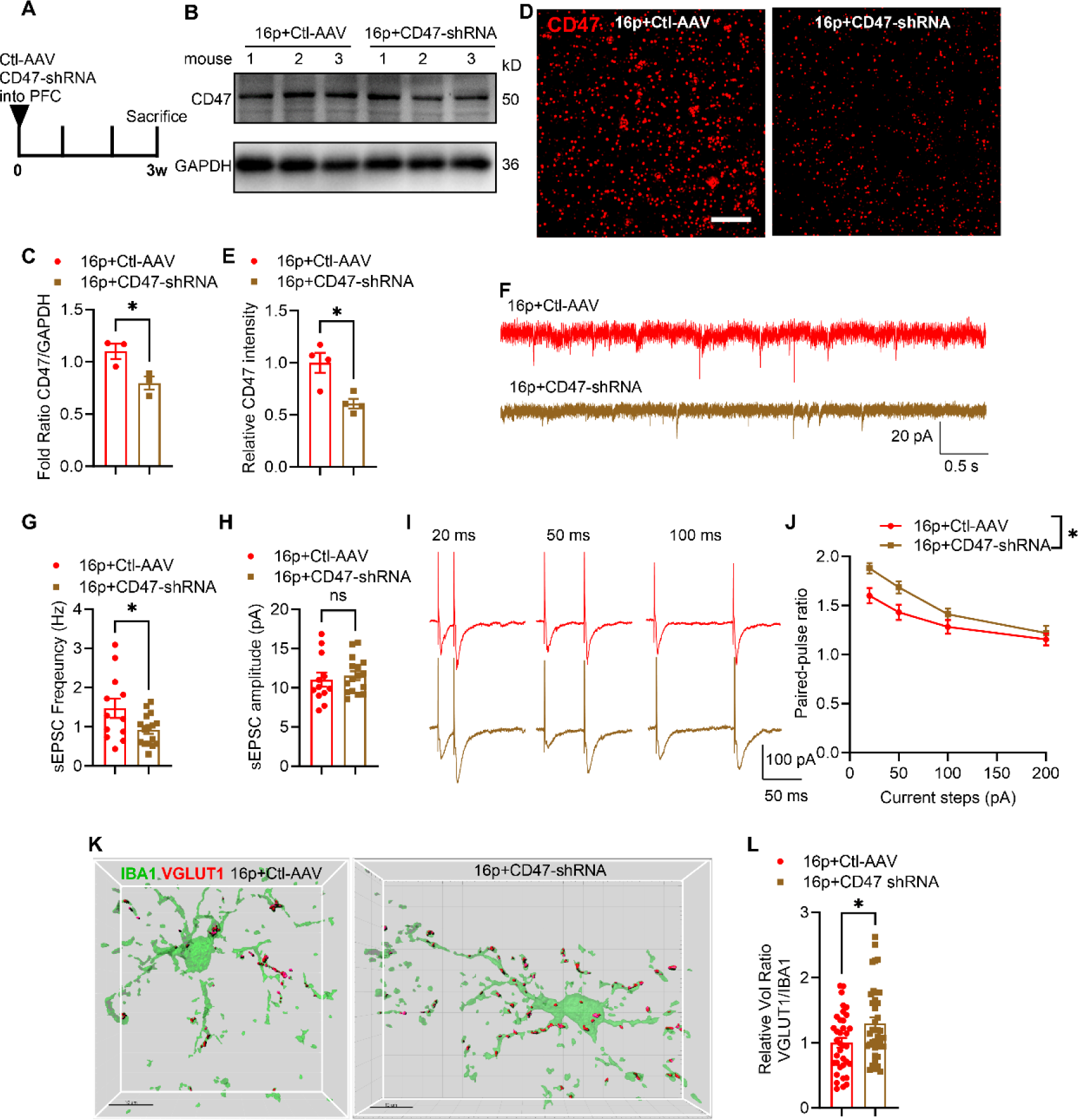
Reducing synaptic CD47 expression *in vivo* reduced excitatory transmission in 16p11.2 deletion mice. A The experimental scheme for stereotaxic injection of control rAAV (Ctl-AAV) and CD47-shRNA in the PFC. B Representative images of CD47 western blotting in two rAAV injected groups. C Quantification of CD47 expression in PFC. D Representative images of PFC CD47 staining in two groups. E Quantification of CD47 intensity in PFC. F Representative traces of sEPSC recorded in PFC in two groups. The scale bar: 20 pA/0.5 s. G Quantification of sEPSC frequency. H Quantification of sEPSC amplitude (16p+Ctl-AAV: n = 12 cells from 4 mice, 16p+CD47-shRNA: n = 16 cells from 4 mice). I Representative traces of paired-pulse stimulation at the intervals of 20 ms, 50 ms, and 100 ms in two groups. The scale bar: 100 pA/50 ms. J Quantification of evoked EPSCs PPR (16p+Ctl-AAV: n = 12 cells from 4 mice, 16p+CD47-shRNA: n = 14 cells from 4 mice). K Representative images of IBA1 and VGLUT1 co-immunostaining in two groups. The scale bar: 10 μm. L Quantification of VGLUT1/IBA1 relative volume ratios in microglia (16p+Ctl-AAV: n = 36 cells from 4 mice, 16p+CD47-shRNA: n = 38 cells from 4 mice). ns, not significant, *P < 0.05, unpaired *t* test for panels C, E, G, H, Mann-Whitney U test for panel L and two-way RM ANOVA with Sidak’s multiple comparisons *post hoc* test for panel J.

### rAAV CD47-shRNA treatment enhanced social novelty ability in 16p11.2 deletion mice

Several behavioral tests were conducted on 16p11.2 deletion mice treated with rAAV CD47-shRNA to assess the potential benefits of modulating CD47. In the open field test, rAAV CD47-shRNA treatment did not significantly affect distance traveled, velocity, and time spent in the center, indicating normal locomotion and anxiety levels (Fig. 6 A–D). The recognition index in the NOR test also remained unchanged, suggesting that rAAV CD47-shRNA treatment did not ameliorate impaired recognition memory (Fig. 6 E and F). In the Y maze test, the alternation index showed no significant alteration either (Fig. 6 G and H). In contrast, the three-chamber test showed that rAAV CD47-shRNA treatment significantly enhanced social novelty ability (Fig. 6 I and K), but did not impact social ability in 16p11.2 deletion mice (Fig. 6 I and J). In summary, reducing synaptic CD47 expression enhanced the social novelty ability of 16p11.2 deletion mice.

**Figure 6.**
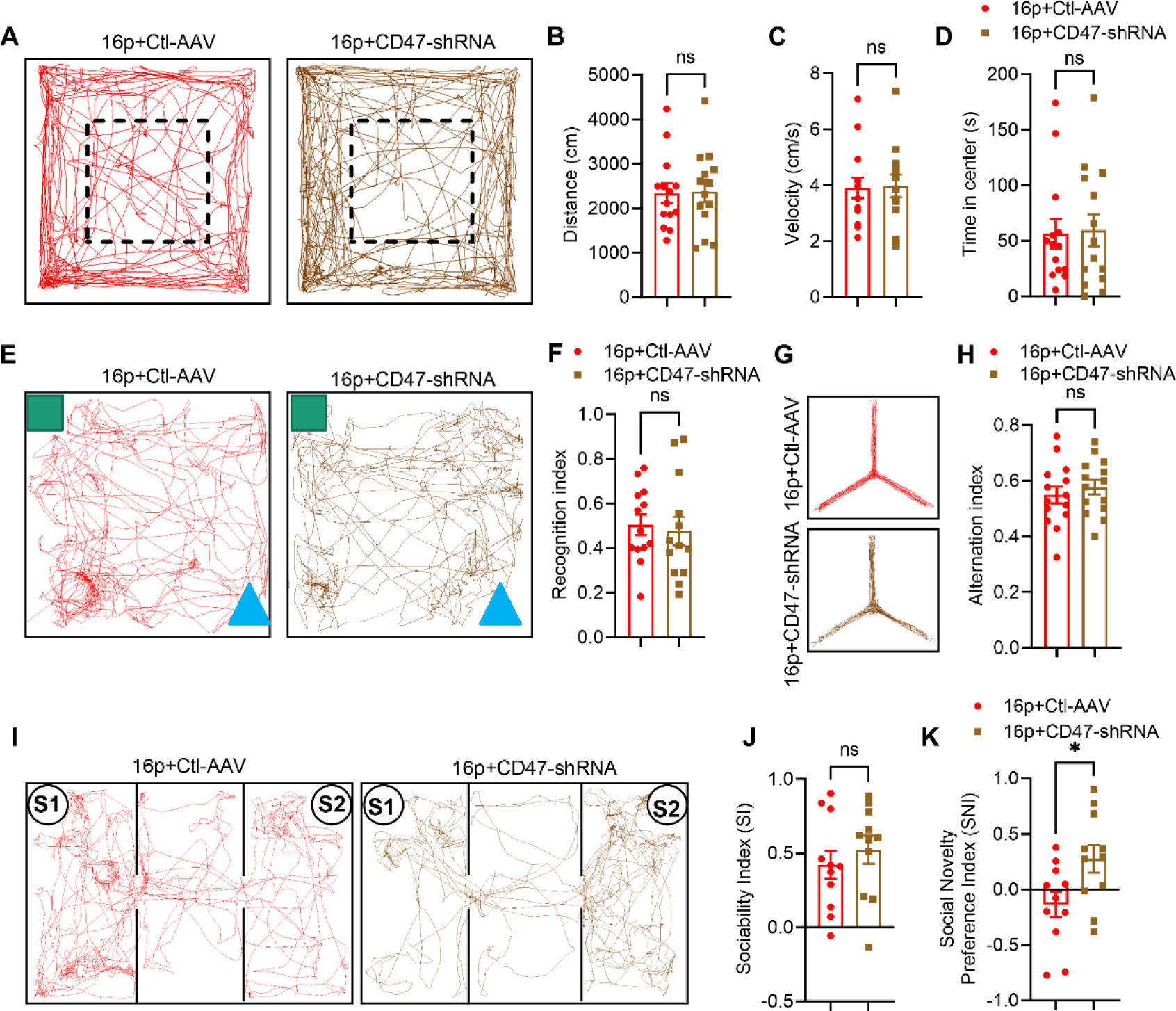
Reducing synaptic CD47 expression enhanced social novelty ability in 16p11.2 deletion mice. A The trajectory chart of mice in the open field. The dash-lined box: center area. B The distances traveled in the open field. C The velocity of movement in the open field. D Time in the center in the open field (16p+Ctl-AAV: n = 14 mice, 16p+CD47-shRNA: n = 14 mice). E The trajectory chart of mice in NOR test. Green square: familiar object; Blue triangle: novel object. F Recognition index in the NOR test (16p+Ctl-AAV: n = 13 mice, 16p+CD47-shRNA: n = 13 mice). G The trajectory chart of mice in Y maze. H Alternation index in the Y maze (16p+Ctl-AAV: n = 14 mice, 16p+CD47-shRNA: n = 14 mice). I The trajectory chart of mice in the three-chamber test. S1: familiar mouse, S2: novel mouse. J Sociability index in the three-chamber test. K Social novelty preference index in three-chamber test (16p+Ctl-AAV: n = 11 mice, 16p+CD47-shRNA: n = 11 mice). ns, not significant, *P < 0.05, unpaired *t* test for panels B, C, F, H, J, K and Mann-Whitney U test for panel D.

## Discussion

The current study revealed a significant link between CD47-mediated impairments in microglial phagocytosis and excitatory transmission and cognitive and behavioral deficits development in the widely used 16p11.2 deletion mouse ASD model. Importantly, the inhibition or reduction of CD47 using a specific antibody or shRNA, respectively, markedly improved microglial synaptic pruning and mitigated social novelty deficits. These findings provide new insights into the mechanism underlying the ineffective synaptic pruning by microglia, which ultimately contributes to defects in excitatory transmission and the manifestation of ASD-like behaviors.

Microglia, resident immune cells in the brain, can phagocytose dead cells and cellular debris during inflammation. Growing evidence indicates that microglia also fulfill a crucial role in the brain’s normal function. Specifically, microglia exhibit a preference for engulfing weaker or less active synapses, thereby facilitating the establishment of well-defined functional neural circuits enriched with stronger or more active synapses. The manifestation of ASD phenotypes has been associated with diminished microglial synaptic pruning (8, 9, 13, 23, 24). However, distinct changes in microglia occur in various ASD animal models. For example, in the prenatal maternal immune activation stress model, some studies have documented an increase in microglia numbers and elevated levels of the proinflammatory cytokine IL-1β (25, 26). In contrast, others show that prenatal immune stress diminishes microglial reactivity due to adverse prenatal conditions, thereby affecting long-term microglial responsiveness and proper striatal circuit development (24). In the present study utilizing the 16p11.2 deletion mouse model, we observed no increase in microglial numbers, surface areas, or volume; however, we noted increased ramification compared to the WT littermates, suggesting a more resting state of microglia. Consequently, 16p11.2 deletion mice exhibited a marked decrease in microglial phagocytic ability, increased synapse number, and excitatory transmission within the PFC. As a result, the 16p11.2 deletion mice display increased local field potential activity in the PFC, diminished long-range prefrontal connectivity (17), and thalamic-prefrontal miswiring and low-frequency neuronal synchronization reduction (21).

Interestingly, the features of microglia in 16p11.2 deletion mice seem to differ from those in the SCN2A deficiency monogenic ASD mouse model (23). The *Scn2a*-deficient mice exhibit decreased synaptic transmission and reduced spine density in hippocampal neurons. In these mice, microglia show partial activation, leading to excessive phagocytic pruning of postsynaptic structures associated with the complement C3 cascades during specific developmental stages. In contrast, in the hippocampal CA1 region of 16p11.2 deletion mice, the collective electrophysiological findings reveal enhanced excitability, disrupted excitation and inhibition balance, and accelerated maturation of glutamatergic synapses (27). Despite these differences, both animal models exhibit impaired learning, memory, and sociability, processes closely linked to the PFC (28) and hippocampus (29). Future studies are required to decipher the selective role of microglia reactivity and synaptic pruning capacity in specific conditions of ASD.

Overexpression of CD47 protects synapses from excess microglia-mediated pruning during development (15) and is associated with brain overgrowth in individuals with 16p11.2 deletion syndrome (18). The current western blotting and immunostaining results for the first time showed that the 16p11.2 deletion mice indeed up-regulated the expression of CD47. CD47 was co-expressed at the synapse, which is positive for VGLUT1 expression and was essential for activity-dependent alterations in engulfment. In CD47 knockout mice, microglia do not prefer to engulf active synapses (14). Thus, the CD47 acts a “don’t eat me” signal overexpressed in the 16p11.2 deletion carriers and contributes to reduced phagocytosis both *in vitro* and *in vivo*.

Based on these observations, CD47 has been suggested as a valid target for treating ASD(18). We tested this hypothesis utilizing a CD47 antibody and rAAV CD47-shRNA. Both interventions were able to reverse deficits in microglial phagocytosis, improve excitatory transmission, and alleviate social novelty deficits in 16p11.2 deletion mice. However, despite these enhancements, impaired recognition memory persisted in these mice following treatment. This lack of impact on the recognition memory could potentially be attributed to the inefficacy of rAAV infection in the PFC. A future molecular genetics approach involving the offspring from crossing CD47 knockout mice with 16p11.2 deletion mice holds promise for elucidating this hypothesis. We anticipate that effective CD47 blockade will ameliorate social novelty and recognition memory deficits in 16p11.2 deletion mice. Nevertheless, the observation that LPS treatment effectively activated microglia and increased microglia phagocytic capacity, as shown in SI Appendix Figure 3, not only confirms the crucial role of microglia in maintaining the homeostatic state of synapses but also suggests that LPS treatment may represent a therapeutic approach for ASD. Indeed, administering a low dose of LPS restored microglial synaptic pruning, normalized synaptic neurotransmission, and rescued behavioral impairments in an ASD mice model lacking Gls1 in CamKIIα-positive neurons (30).

Our study underscores the significance of neuronal CD47 in facilitating microglial synaptic pruning, thereby improving excitatory transmission and mitigating behavioral deficits in 16p11.2 deletion mice. This study represents the first demonstration that targeting CD47 holds promise as a therapeutic avenue for enhancing outcomes in ASD. Further investigations are warranted to consolidate the role of CD47 interference as an efficacious therapeutic strategy for ASD.

A limitation of this mouse model is that it only reflects a small proportion (5.5%) of autism cases linked to genetic factors (3). To determine the broader implications of CD47 knockdown, future studies should investigate its effects on more prevalent murine models of autism, such as the BTBR mice model (31) and the prenatal maternal immune stress models like the VPA and MIA models (25, 32). The comparative examination will be pivotal in gauging the clinical significance of CD47 intervention as a prospective therapy for ASD.

## Materials and Methods

### Animals

Mice carrying the 16p11.2 deletion on a hybrid C57BL/6N x 129Sv genetic background were obtained from Jackson Laboratory and bred locally (JAX:013128). Both male and female mice with the 16p11.2 deletion aged 2-3 months were selected for the study. Mice were housed in a controlled environment in a pathogen-free SPFII animal facility at 23 ± 1 °C and 50 ± 10% humidity and were subjected to a 12:12 h light/dark cycle (7 a.m. to 7 p.m.) with a light intensity of 15 - 20 lx during the light period, except during cleaning and experimental procedures when the room light was at 200 lx. Mice were group-housed in ventilated cages with six animals per cage and provided food and water *ad libitum*. Experimenters remained blinded to the animals’ treatments and sample processing throughout the experimentation and analysis.

### Ethical approval and animal experimentation design

The Animal Care Committee of the Southern University of Science and Technology (Shenzhen, China) approved the animal experiment protocols. The ARRIVE guidelines for designing, performing, and reporting animal experimentation were followed (33). Mice in the study were randomly assigned to groups to ensure total randomization. Efforts were made to minimize animal numbers and suffering. Inclusion criteria were based on the identical age and sex of the mice. The AEEC Animal Experimentation Sample Size Calculator was utilized to determine the minimum sample size needed for the study hypothesis (34). Results indicated that a minimum of six mice per group were required for behavioral studies to achieve meaningful statistical differences. Additionally, at least three mice per group were used for slice electrophysiology, immunostaining, and the Golgi staining.

### Mouse behavioral tests

#### Open field test

The open-field test was conducted to evaluate locomotion and anxiety levels in mice, following a previously described method (35). Prior to the test, mice were allowed to acclimate to the testing room for one hour. Each mouse was then placed in the center zone of the open field (40 x 40 x 40 cm), which was divided into 16 sections, with the four middle sections (20 cm x 20 cm) designated as the center area. The EthoVision XT software from Noldus Information Technology (Leesburg, USA) was used to record the total distance traveled by the mice and the time spent in the center area during a 10-minute period.

#### Y maze test

The Y maze test was conducted to assess spontaneous alternations, reflecting working memory capacity. The maze consists of three arms with high walls and a triangular central platform (30 cm x 5 cm x 15 cm), each arm featuring visual cues at the end. Prior to the test, mice were given 1 hour to explore the testing room. During the test, each mouse was placed on the central platform and allowed to move freely for 5 minutes. The sequence and total number of arms entered by each mouse were recorded using EthoVision XT software. Percentage alternations were then calculated using the formula: percentage alternations = number of three consecutive entries into a new arm / (total number of arms entered - 2).

#### Novel object recognition test

The experiments were carried out in an open field box (40 cm × 40 cm × 40 cm) as previously described (36). Prior to testing, the mice were allowed to acclimate to the test room for one1 hour. In the initial stage, two identical plastic toys were placed in the corners of the box, and the mice had 10 minutes to explore them. Subsequently, the mice were removed from the box and returned to their home cage. Two hours later, the mice were reintroduced to the test box, with one of the plastic toys replaced by a new toy of similar size but different color and shape. The mice were given 10 minutes to explore these objects. The time spent by the mouse’s nose tip within a statistical range of 6 cm x 6 cm was measured, with exploration time automatically recorded by the EthoVision XT software.

#### Three-chamber test

The method used were as we previously described. The social behavior of mice was studied using a three-chamber apparatus that was divided into three interconnected chambers with transparent plexiglass. The mice were first habituated to the apparatus for 10 minutes. Sociability was then evaluated during a second 10-minute period, during which the test mice could interact with either an empty cage or a genotype, age, and sex-matched stranger mouse (Mouse 1) that was placed in a cage in one of the chambers. Preference for social novelty was then assayed in a third 10-minute period by introducing a second stranger mouse (Mouse 2) into the previously empty cage. The time spent interacting with the empty cage, Mouse 1, or Mouse 2 was recorded, and the statistical range of 14 cm x 14 cm was measured using EthoVision XT 10 software. The sociability index (SI) and the social novelty preference index (SNI) were calculated as follows:

Sociability index:

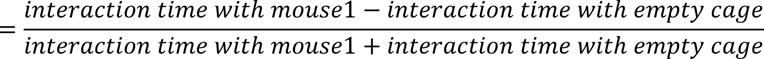

Social novelty preference index:

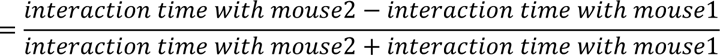

### Stereotaxic surgery

The methods used were as we described previously (37). The mice were anesthetized with isoflurane and then received bilateral injections of 250 nanoliters of AAV2/9-Syn-shRNA CD47-ZsGreen (concentration at 1.9E+12 vg/ml; vg, viral genome) or AAV2/9-Syn-ZsGreen (concentration at 1.7E+12 vg/ml) from HANBIO Technology (China), into the PFC at a rate of 0.05 microliters per minute. The injection site was targeted at AP: +1.98 mm, ML: ±0.3 mm, DV: −2.0 mm from bregma. The needle remained in place for 5 minutes post-injection before being removed. The mice were allowed to recover in their home cage until fully awake. To reduce pain, meloxicam (1 mg/kg, s.c.) and penicillin (3000 U per mouse, i.p.) were administered once daily for 3 days. The AAV virus was expressed for three weeks to label neurons with ZsGreen. The nucleotide sequence of the shRNA CD47 is 5’-CACCGAAGAAATGTTTGTGAA-3’.

### Brain slice electrophysiology for sEPSC, PPR, and AMPA/NMDA ratio

The protocol for brain slice preparation was adapted from previous studies (36–38). Mice were anesthetized with 1% pentobarbital sodium and euthanized by decapitation. The mouse brain was dissected and immersed in ice-cold artificial cerebrospinal fluid (aCSF) containing (in mM): 30 NaCl, 26 NaHCO_3_, 10 D-glucose, 4.5 KCl, 1.2 NaH_2_PO_4_, 1 MgCl_2_, and 194 sucrose. Additionally, 1.5 mL of 1M HCl per 1 L of cutting solution was added, and the solution was bubbled with 95% O_2_/5% CO_2_. Coronal brain slices (350 μm) were prepared using a vibratome (VT1120S, Leica Systems, Germany). The slices were allowed to recover for 30 minutes at 34°C in aCSF containing (in mM): 124 NaCl, 26 NaHCO_3_, 10 D-glucose, 4.5 KCl, 1.2 NaH_2_PO_4_, 1 MgCl_2_, and 2 CaCl_2_. Additionally, 10 g of sucrose and 1 mL of 1M HCl per 1 L of aCSF were added, and the solution was bubbled with 95% O_2_/5% CO_2_. Following transfer to a holding chamber at room temperature, recordings commenced only after at least 1 hour of recovery. The slices were then positioned in a recording chamber (RC26G, Warner Instruments, USA) on the x-y stage of an upright microscope (BX51W; Olympus, Japan) and perfused with aCSF at a 2 ml/min rate. All recordings were conducted at room temperature.

The sEPSCs, PPR, and AMPA/NMDA ratio in pyramidal neurons were recorded using patch clamping techniques. The recording pipettes were filled with the following solution (in mM): 125 CsMeSO_3_, 5 NaCl, 10 HEPES (Na^+^ salt), 5 QX314, 1.1 EGTA, 4 ATP (Mg^2+^ salt), and 0.3 GTP (Na^+^ salt). sEPSCs of pyramidal neurons located in the PFC were recorded in aCSF supplemented with 20 μM bicuculline at a holding potential of −60 mV. The PPR was recorded to evaluate the probability of presynaptic glutamate release. Electrical stimulation (0.1 ms square pulse) was administered using a glass electrode filled with aCSF, positioned within 0.1 mm of the recording site. Throughout the recording session, aCSF containing 20 μM bicuculline was continuously perfused. PPR recordings were conducted using two stimulations with durations of 20 ms, 50 ms, 100 ms, and 200 ms. To obtain the NMDAR–EPSC-to-AMPAR– EPSC ratio (AMPA/NMDA ratio), AMPAR–EPSC was first recorded at −60 mV. Then, the mixture of AMPAR–EPSC and NMDAR– EPSC was recorded at +40 mV with the same stimulation pulse (0.1 ms). The peak of NMDAR– EPSC was calculated at 50 ms from the onset of the EPSC mixture. Throughout the recording session, aCSF containing 20 μM bicuculline was continuously perfused. The acquisition frequency was set at 20.0 kHz, and the filter was adjusted to 2.9 kHz. sEPSCs were analyzed using Mini-analysis (Synaptosoft Inc.), focusing on frequency and amplitude. Additionally, PPR, AMPAR– EPSC and NMDAR–EPSC traces were imported into Fitmaster software (HEKA Elektronik) to measure evoked EPSC amplitude.

### Immunofluorescence staining

Mice were anesthetized with phenobarbital sodium salt (0.1 g/kg) and perfused transcardially with ice-cold 0.01 M PBS, followed by 4% paraformaldehyde (PFA) in 0.01 M PBS. The brains were fixed and cryopreserved in 4% PFA overnight, then dehydrated in 30% sucrose. Coronal sections (30 μm) were prepared and blocked with a solution containing PBS, 0.3% Triton X-100, and 10% goat serum for 1 hour at room temperature. The sections were then incubated overnight at 4°C with primary antibodies: anti-IBA1 (rabbit, 1:500, ab178847, Abcam), anti-VGLUT1 (guinea pig, 1:500, 135304, Synaptic Systems), anti-CD68 (rat, 1:500, MCA1957, Bio-Rad), anti-PV (rabbit, 1:1000, ab11427, Abcam), anti-NeuN (mouse, 1:500, ab104224, Abcam), anti-CD47 (rabbit, 1:200, ab214453, Abcam). After primary antibody incubation, sections were treated with appropriate secondary antibodies for 1 hour at room temperature. Visualization was performed using a Zeiss LSM980 confocal microscope.

### Analysis of 3D microglia engulfment and microglia morphology

The method used to analyze microglial morphology was precisely as we previously described. The z-stack images (at 1 μm intervals) were captured using a Zeiss LSM980 confocal microscope with a 63X lens. Microglia were selected randomly based on positive staining for IBA1, ensuring an unbiased approach. Individual microglia, including those with VGLUT1 signaling, were isolated using Image J and transferred to Imaris 9.0.0 software (Oxford Instruments, USA). Initially, 3D volume surface renderings of the microglial channels were generated to compute each microglial cell’s volume and surface area. Subsequently, a new channel representing engulfed VGLUT1 was established using the mask function to determine the volume of engulfed VGLUT1. The engulfment percentage was calculated as the volume of internalized VGLUT1 puncta divided by the volume of the microglial cell (15, 39). Next, using the filament tool, individual microglia were traced using the default settings (autopath (no loops) and cell body sphere region ∼10 μm) and manually modified if necessary, and filament dendrite length of microglia was measured automatically (40).

### The Golgi staining

The Golgi staining was conducted using the FD Rapid GolgiStainTM kit (FD NeuroTechnologies, Ellicott City, MD, USA) following the manufacturer’s instructions and as previously described (41). Briefly, the brains were rinsed with double distilled water (ddH_2_O) and then immersed in a 1:1 mixture of FD Impregnation Solution A and B in darkness for 3 weeks at room temperature. The solution was replaced once every 24 hours. Following impregnation, the brains were transferred to FD Solution C and stored in darkness for 5 days, with the solution being replaced after 24 hours.

The individual brains were mounted on a specimen disc with optimum cutting temperature compound and subjected to snap freezing and cryosectioning on a Leica CM1950. Coronal sections of 200 µm thickness were then cut and transferred to agar-treated slides for staining. After drying for 4 days, the brain sections were rinsed twice with ddH_2_O and stained in a mixture of FD Solution D, FD Solution E, and ddH_2_O in a 1:1:2 v/v ratio for 10 minutes. The stained sections were rinsed twice with ddH_2_O for 4 minutes each and sequentially dehydrated in 50%, 75%, 95%, and 4 times in 100% ethanol, with each dehydration step lasting 4 minutes. Following dehydration, the sections were cleared 3 times in xylene for 4 minutes each rinse and sealed with a resinous mounting medium. The Golgi-stained sections were examined under an optical microscope at 100X magnification, and the images were analyzed using Image J software to determine the density per 10 µm of dendritic length.

### Western blotting

The method used was essentially as we previously described (38). Briefly, the hippocampus was removed and lysed in 200 ml lysis buffer (Cat. #C500008, Sangon Biotech, China) containing proteinase and phosphatase inhibitors. The lysates were centrifuged at 13,680 g for 10 min at 4 °C, and the supernatant was collected and boiled in boiling water for 10 min with a loading buffer (4:1 ratio). Whole proteins were electrophoresed in a 10% SDS-PAGE gel and then transferred to polyvinylidene fluoride (PVDF) membranes (0.45 μm) using iBlot 2 Dry Blotting System (Thermo Fisher Scientific Systems, USA). After blocking in 5% skim milk, the PVDF membranes were incubated with the primary antibody (anti-IBA1: rabbit, 1:1000, ab178847; anti-CD47: rabbit, 1:1000, ab214453; anti-CORO1A, rabbit, 1:1000, ab228635; anti-GAPDH, rabbit, 1:3000, ab181602; anti-β-ACTIN, mouse, 1:3000, ab6276) overnight at 4 °C. The PVDF membranes were then incubated with a secondary antibody after washing three times with TBST. Protein band intensities were detected with Tanon (Shanghai, China).

### Statistical Analysis

All data were expressed as the Mean ± Standard Error of the Mean (SEM). Statistical analyses were conducted using Prism (V9, GraphPad Software, USA). Data distribution was assessed initially using the Shapiro-Wilk test to determine suitability for parametric or nonparametric tests. An unpaired *t*-test was employed for normally distributed data in two-group comparisons. Non-normally distributed data were analyzed using the Mann-Whitney U test. The paired-pulse ratio was evaluated using two-way repeated measures ANOVA with Sidak’s *post hoc* tests. Specific experiment details, including exact sample sizes (n), precision measures, statistical tests performed, and definitions of significance, are provided in figure legends. Statistical significance was set at P < 0.05.

## Data Accessibility

All primary data are archived at the Southern University of Science and Technology and available upon request.

## Acknowledgments

We express our gratitude to the SUSTech Animal Facility for their assistance with animal maintenance and to the SUSTech Core Research Facilities for their support with imaging acquisition. Financial assistance for Jun Ju was received from the National Natural Science Foundation of China (82301730), and Dr. Ju is a Pengcheng Peacock Plan (C) Distinguished Research Professor. Financial support for Dr. Sheng-Tao Hou was provided through grants from the National Natural Science Foundation of China (32371029), the Shenzhen-Hong Kong Institute of Brain Science-Shenzhen Fundamental Research Institutions (2023SHIBS0002), and the Shenzhen Medical Research Fund (B2301001). Dr. Hou also receives support from the Guangdong Innovation Platform of Translational Research for Cerebrovascular Diseases and the SUSTech-UQ Joint Centre for Neuroscience and Neural Engineering (CNNE). Dr. Hou holds the title of Pengcheng Peacock Plan (A) Distinguished Professor.

