## Supplementary material for "The “don’t eat me” signal CD47 contributes to microglial phagocytosis defects and autism-like behaviors in 16p11.2 deletion mice": Supplemental Files.pdf

**This PDF file includes:**

Supporting text

Figures S1 to S5

Legends for Figures S1 to S5

**Supporting Information Text**

### SI Appendix Figure 1

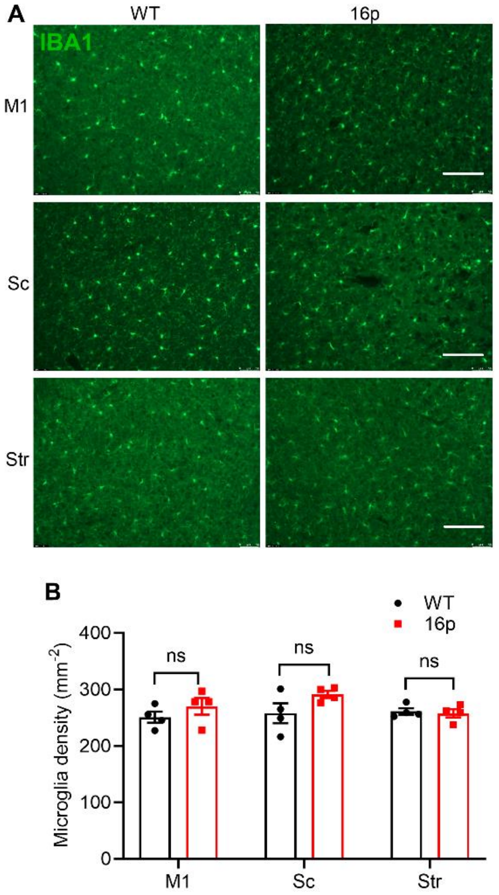


**Supplementary Figure 1. Unaltered microglial numbers in different brain regions in 16p11.2 deletion mice**

A Representative image of IBA1 microglia immunostaining in the primary motor cortex (M1), sensory cortex (Sc), and striatum (Str) in two mice groups. The scale bar: 100 µm. B Quantification of microglia intensity. The number of mice: WT n = 4 mice, 16p n = 4 mice; ns, not significant, unpaired *t* test.

### SI Appendix Figure 2

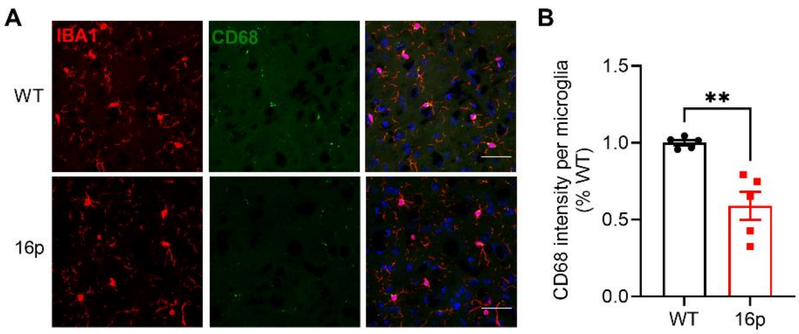


**Supplementary Figure 2. Reduced CD68 expression in microglia in 16p11.2 deletion mouse PFC**

A Representative images of IBA1 and CD68 co-immunostaining in two groups of mice. The scale bar: 100 μm. B Quantification of CD68 intensity per microglia (WT: n = 5 mice, 16p: n = 5 mice). **$P < 0.01$, unpaired *t* test.

### SI Appendix Figure 3

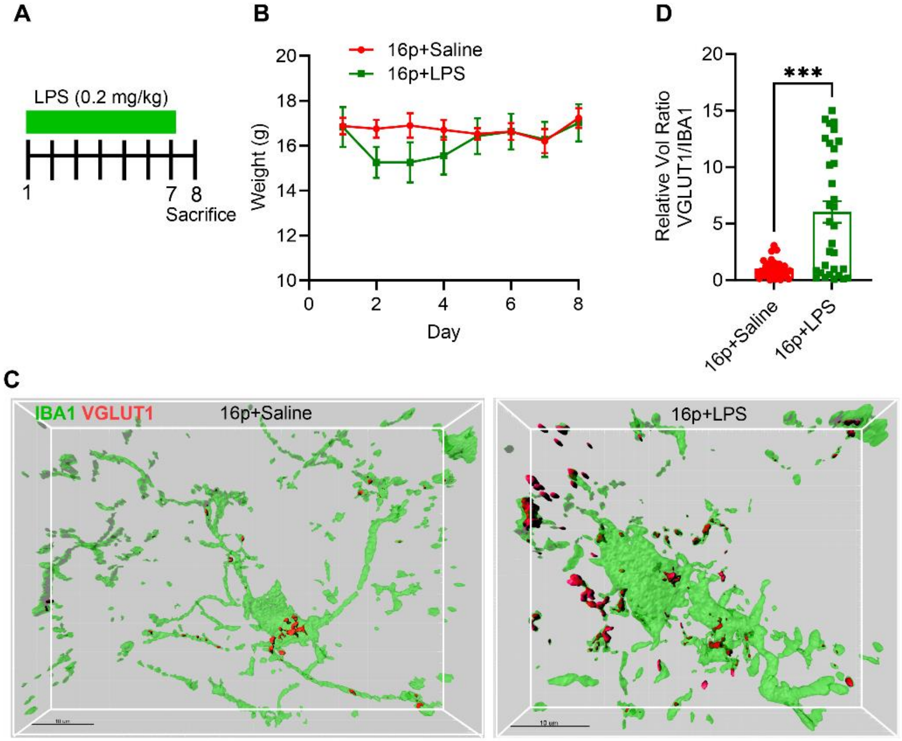


**Supplementary Figure 3. Activation of microglia using LPS promoted microglia-dependent synapse pruning in 16p11.2 deletion mice**

A The experimental scheme of LPS treatment. B LPS transiently reduced body weight in 16p11.2 deletion mice (16p+saline: n = 4 mice, 16p+LPS: n = 4 mice in each group). C Representative images of IBA1 and VGLUT1 co-immunostaining in two groups of mice. The scale bar: 10 μm. D Quantification of VGLUT1/IBA1 relative volume ratios in microglia (16p+saline: n = 38 cells from mice, 16p+LPS: n = 32 cells from 4 mice). ***P < 0.001, Mann-Whitney U test.

### SI Appendix Figure 4

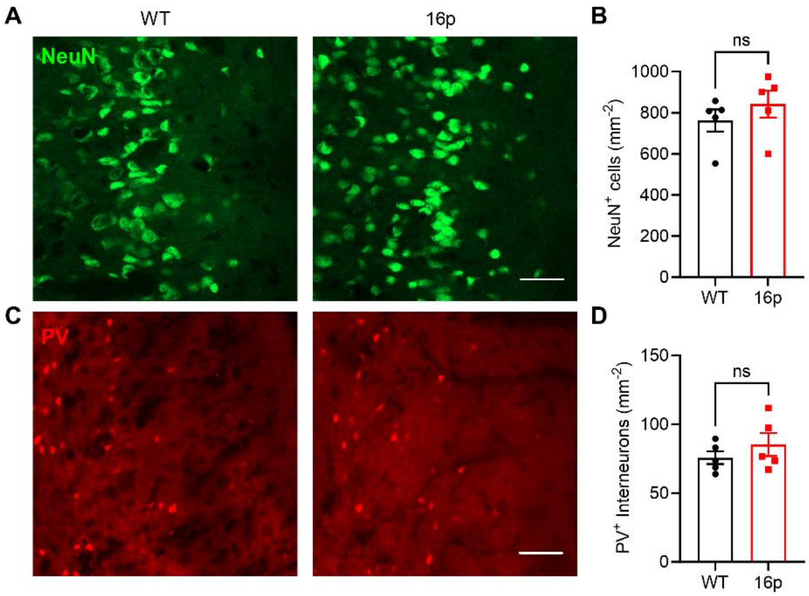


**Supplementary Figure 4. Unaltered numbers of mature neurons and PV interneurons in 16p11.2 deletion mouse PFC**

A Representative images of PFC immunostained with a NeuN antibody in two groups of mice. The scale bar: 100 μm. B Quantification of the density of $NeuN^+$ cells in the PFC (WT: n = 5 mice, 16p: n = 5 mice). C Representative image of PV interneurons in two groups of mice. The scale bar: 100 μm. D Quantification of the density of $PV^+$ interneurons in the PFC (WT: n = 5 mice, 16p: n = 5 mice). ns, not significant, unpaired *t* test.

### SI Appendix Figure 5

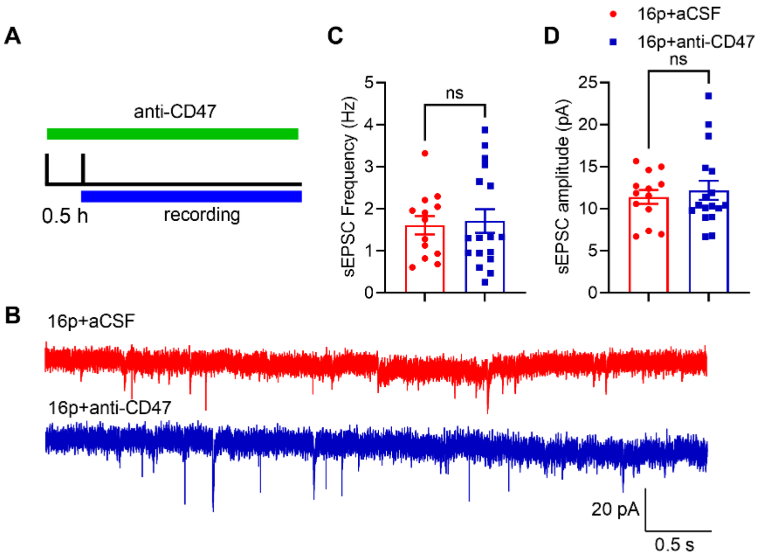


**Supplementary Figure 5. Blocking PFC CD47 using a specific antibody did not affect excitatory transmission in 16p11.2 deletion mice brain slice**

A The experimental scheme. B Representative traces of sEPSC recorded in PFC in two groups of mice. The scale bar: 20 pA/0.5 s. C Quantification of sEPSC frequency. D Quantification of sEPSC amplitude (16p+aCSF: n = 13 cells from 3 mice, 16p+anti-CD47: n = 17 cells from 3 mice). ns, not significant, Mann-Whitney U test.
